# The distribution of functional N-cycle related genes and nitrogen in soil profiles fertilized with mineral and organic N fertilizer

**DOI:** 10.1101/2020.01.15.907543

**Authors:** Massimo Zilio, Silvia Motta, Fulvia Tambone, Barbara Scaglia, Gabriele Boccasile, Andrea Squartini, Fabrizio Adani

## Abstract

Nitrogen (N) fertilizers applied to agricultural soils result in the release of nitrogen, mainly nitrate (NO_3_^-^) in addition to nitrous oxide (N_2_O) and ammonia (NH_3_), into the environment. Nitrogen transformation in soil is a complex process and the soil microbial population can regulate the potential for N mineralization, nitrification and denitrification. Here we show that agricultural soils under standard agricultural N-management are consistently characterized by a high presence of gene copies for some of the key biological activities related to the N-cycle. This led to a strong NO_3_^-^ reduction (75%) passing from the soil surface (15.38 ± 11.36 g N-NO_3_ kg^-1^ on average) to 1 m deep layer (3.92 ± 4.42 g N-NO_3_ kg^-1^ on average), and ensured low nitrate presence in the deepest layer. Under these circumstances the other soil properties play a minor role in reducing soil nitrate presence in soil. However, with excessive N fertilization, the abundance of bacterial gene copies is not sufficient to explain N leaching in soil and other factors, i.e. soil texture and rainfall, become more important in controlling these aspects.

## Introduction

Anthropogenic activities are the major driver of changes in the global nitrogen (N) cycle since the last century, resulting in N-flows being 3.3-fold higher than those due to natural processes, achieving a total globally fixed nitrogen of 413 Tg N y^-1^. Since nitrogen is one of the most important nutrients for many life forms, such a strong change in its availability has important effects on the balance of terrestrial and aquatic ecosystems [1,2].

Agriculture has indeed a major role in this process: the global amount of N used in agriculture has increased from 12 Tg N in 1960 to 104 Tg N in 2010 and the amount of N_2_ fixed to NH_4_^+^ by industrial processes and destined for agriculture contributes today to 45% of the total nitrogen fixed annually on the planet [1,3,4].

As a consequence of that, the total amount of N brought to the soil today, on a world scale, is more than twice that considered a safe planetary boundary, i.e. a safe operating space for humanity to avoid the risk of ecosystems’ deterioration [5–7]. Furthermore, an increase in the use of N in the food system by 51% in 2050 has been estimated, with a global increase in the environmental pressure of the food system estimated to be in the range 52%-90%, in the absence of mitigation measures [7,8]. Without emission reductions, global N losses are expected to further increase, reaching in 2050 levels equal to 150% of those of 2010 [9].

Nitrogen fertilizers applied to agricultural soils result in the release of N into the environment, both in the atmosphere (NH_3_, N_2_O, N_2_), and in groundwater (NO_3_^-^) [10]. In many areas of the world, agriculture has been acknowledged as the single largest source of N (mainly NO_3_^-^) to environments [11–14], causing alterations in the carbon cycle, biodiversity reduction, acidification, soil fertility reduction and air pollution [15]. Furthermore, human health problems should be also considered; for example, in recent years many studies reported an increased incidence of colorectal cancer in subjects who regularly drink water with a concentration of nitrate above 8.6 mg L^-1^ [16,17]. All these problems lead to a social cost that has been estimated to be up to 10 $ kg^-1^ N [18].

Nitrogen in soils can be metabolized and transformed by soil microbial communities [3,19–23], returning to the atmosphere in gaseous form (N_2_O, N_2_). Once the reduced forms of ammonia, coming either from mineral or organic fertilization, are oxidized by nitrifiers, the denitrification process can ensue which reduces nitrate and nitrite (NO_2_^-^) to nitric oxide (NO), nitrous oxide (N_2_O), and dinitrogen (N_2_).

Today there is general agreement that soil microorganisms play a central role in the N cycle [3,20,22] as they are responsible for the conversion of N in its various forms: the structure of soil microbial populations is regarded as the major variable that regulates the potential for N fixation, mineralization, nitrification and denitrification [24].

In the recent years some findings have enriched the complexity of the known pathways ruling the nitrogen cycling in the environment. One of such cases is the comammox [25], in which one single species of the genus Nitrospira has been demonstrated to perform a complete phenotype of nitrification from ammonium to nitrate. Another previously overlooked distinction is the split of the nitrous oxide reductase genotype in two clades (*nosZ* I and *nosZ* II) the second of which corresponds to novel non-denitrifying types of N_2_O reducers [26].

Notwithstanding the existence of other genetic determinants linked to the N cycle that, as just commented, can occur in soils in the subset of key nitrogen cycle genes (archaeal *amoA*, eubacterial *amoA, nirK, nosZ* and *nifH*) that we chose to include as suitable proxies for nitrification, denitrification and nitrogen fixation in our analysis was defined based on the following considerations. With respect to the comammox *Nitrospira*, which was originally discovered upon enriching cultures from material found in a biofilm growing within a steel pipe in deep oil wells [25] subsequent studies had addressed the relevance of such taxa in agricultural contexts [27] and concluded that although the species is present in soils, the dominant contributors of potential nitrification are the classic ammonia oxidizing bacteria and the newly discovered comammox do not play a significant role in these pathways (P < 0.05).

Concerning the second *nosZ* clade (*nosZ* II) we chose to restrict the survey of the terminal gene of denitrification to *nosZ* I based on the conclusions of Domeignoz-Horta and coworkers (2015) [28] who reported that: (a) the *nosZ* I community was consistently more abundant than the *nosZ* II one and (b) no significant differences between the two groups could be ascribed to the different agricultural management practices, neither in relation to crops nor to fertilization regimes. The same authors add that the lack of detectable variations between these subgroups is in line with the fact that such differences have been reported only in long-term agronomical trials that had been carried out for over 50 years.

Finally, regarding the known existence of two families of genes able to perform nitrite reductase activity converting nitrite into nitrous oxide (*nirK* and *nirS*) we selected the former due to the following reasons: (a) *nirK*-harboring bacteria mostly dominate in soils and rhizospheres over nitrite reducers of the nirS kind [29]; (b) there is a tight correlation between *nirS* and *nosZ* [30] which allows to infer indirect information on the abundance of the former by analyzing the latter. These considerations are also confirmed by our prior work [31] in which we analyzed both *nirK* and *nirS* as well as *nosZ* in Bermuda grass rhizospheres.

Enhanced N removal through optimization of denitrification has drawn much attention as an effective approach towards N control because it is the only pathway, except for the process of anaerobic ammonium oxidation (Anammox), by which reactive forms of nitrogen (Nr) in terrestrial and aquatic ecosystems are transformed back into inert N_2_ gas [19,32]. When not converted into gaseous forms, N stored in soil can be progressively leached as NO_3_^-^ polluting groundwater and shallow water bodies [33–35]. Understanding the dynamics of N transformation and movement in soils is complex because of the large number of variables affecting this process [19,33,36–39].

Nitrogen fertilization and the type of N fertilizers have also been reported to affect N-related microbial populations [40]. Indeed, a drastic effect on the balance ammonia oxidizing archaea (AOA) vs. eubacteria (AOB) was observed, with the AOBs stimulated by the providing of N to the soil, especially organic N, while the AOBs seems to be less responsive, or even inhibited by both mineral and organic fertilization [41]. For denitrifying bacterial populations, a generic increase in their number was reported in long-time N-fertilized soils compared to unfertilized, especially in the case of organic fertilizer use (manures). On the other hand, fertilizations with sewage sludge showed a negative effect on these microorganisms, probably due to the acid pH of this type of fertilizer [42].

In addition, environmental factors, such as pH, soil texture, humidity and nitrogen availability, are reported to affect the structure and size of soil microbial populations, though the interdependency of these factors with microbial populations in relation to the N-cycle is still largely unclear [22,43–48].

This study aims at determining the potential of agricultural soils to reduce nitrate concentration-down to one-meter depth, focusing on the role of soil microorganisms related to the N-cycle in transforming reactive N. To do this, we investigated reactive forms of nitrogen distributed along the profiles (0-25 cm, 25-50 cm, 50-75 cm and 75-100 cm) of ten different agricultural soils located in the Po Valley (northern Italy), one of the most intensive agricultural areas of the EU, managed with different N-fertilization for both total N dosed and N-fertilizers type, during three growing seasons. The N cycle-related microbial communities were PCR-quantified down the soil profiles [49], and data collected from 308 samples were critically compared with chemical data and soil properties to draw a clear picture of the potential of soil in reducing nitrate-leaching.

## Material and Methods

### Experimental sites

Twelve soil sites cultivated with cereals (mainly corn) and distributed in eight different localities in the Po valley (Italy) (S1 Fig) were considered in the years 2014, 2015, 2016. Ten of them (soil codes 1a, 1b, 2, 3a, 3b, 4a, 4b, 5, 6a, 6b) were fertilized by a regular farming approach using different types and quantities of nitrogen up to a maximum of 450 kg N ha^-1^ (Stage 1 of the study). The last two soils (soil codes 7 and 8) received in 2016 an excess of N fertilizers (Stage 2 of the study). In particular, Soil 7 was equivalent to Soil 4a (S4 Table) but it received an extra N-fertilization in October (860 kg N Ha^-1^, for a total annual N of 1,243 kg N Ha^-1^) by using pig slurry (S1 Table). Soil 8 (S4 Table) represented a field cropped with maize and receiving, during the season an excess of N (1,470 kg N Ha^-1^) by three N-fertilization events during the year, using digestate in April, urea at the end of July and pig slurry in October (S1 Table). Soils with the same code number but different letters were carried out at the same site (farm), but in different fields and with different N fertilizers. The agronomic management of each site is reported in S1 Table.

### Sampling

For each experiment, between March 2014-October 2016, soil cores at 4 depths (0-25 cm; 25-50 cm; 50-75 cm; 75-100 cm) were collected. At least 5 samples were taken for each experiment during the crop cycle, in correspondence with the principal phenological plant stages. Sampling periods are reported in S2 Table.

For each soil sampling a composite sample was taken, formed by mixing 10 sub-samples taken inside the plot. The collection points within each experimental plot were identified according to an X distribution, taking care to avoid the borders of the plots.

Soils taken for chemical analyses were put into sealed containers and stored at 4°C; analyses were performed starting the next day. Soil samples for DNA extraction and qPCR were processed in the hours immediately following the sampling.

### DNA extraction and quantification

Total DNA extractions were carried out on a quantity of soil equal to 5g per sample. For total DNA extraction, the NucleoSpin® Soil (Macherey-Nagel) kit was used. The total DNA extracted was then quantified and normalized using the Quant-iT™ PicoGreen™ dsDNA Assay Kit (Thermo Fisher Scientific). Real time PCR reactions were performed on real-time 7900HT (Applied Biosystems) using SyberGreen technology, in a final volume of 10μl. The sequences of the primers used are reported in S3 Table. Each sample was tested in triplicate, and the standard calibration curve was built using five points in triplicate, equal to fifteen reactions. As templates for the standard curves, amplicons for each of the target genes were cloned into purified plasmids (pGem-T; Promega Corp.) and inserted into *E. coli* JM101 by electroporation. Knowing the size of the vector (3,015 bp) and those for each insert (data from literature, S3 Table), and measuring the plasmid DNA concentration, the number of copies per ng of DNA and the corresponding amounts to be used for each of the quantitative PCR calibration curves, were calculated. Data analysis was performed using SDS v2.1 software (Applied Biosystems).

### Nitrogen and phosphorus analysis

The soil cores were frozen after collection and analyzed for the chemical characterization, and then the soil nitrate concentration (N-NO_3_) was determined according to ISO 13395 (1996), the soil ammonia content (N-NH_4_) according to ISO 11732 (1997) and the soil phosphorus content (P_2_O_5_) according to Watanabe and Olsen (1965). All the analyses were performed in triplicate. The values shown represent the average of the three replicates, with a standard deviation always below 5%.

### Soil parameters analysis

For each site, a soil profile analysis was performed in order to determine the main soil layers and their physical and chemical characteristics. The main soil parameters were determined as follow: soil pH in aqueous solution using a 1:2.5 sample/ water ratio, total organic carbon by dichromate method. For cation exchange capacity (CEC) determination samples were saturated with BaCl_2_-triethanolamine solution (pH 8.1).

### Statistical analysis

Principal Component Analysis (PCA) was performed using the excel package Addinsoft XLSTAT™. For two-way ANOVA analysis the IBM SPSS™ Statistics software was used, with a significance threshold set at 0.05. Multiple comparisons were performed with the Duncan and Gabriel methods. When necessary, datasets were normalized as log_10_x, and homogeneity of variances was tested through the Levene test.

Multiple linear regressions for nitrate content in soils (75-100 cm) (*n* = 10) vs. agronomical, chemical, physical, biological and meteorological data (parameters = 26) were done using the partial least square method (PLS). The cross-validation leave-one-out approach of un-scaled variables was applied to calculate the goodness of regressions (goodness of fit coefficient-R^2^ and goodness of prediction coefficient-R^2^cv, respectively). Taking into consideration all variable values, the PLS regression was calculated and the importance of each independent variable (importance coefficient) defined. Then PLS analysis was repeated removing those variables characterized by the coefficient of the least importance. This procedure was repeated until a final regression model with goodness of regressions coefficient (R^2^ and R^2^cv) and the smallest number of variables was achieved. PLS was performed using SCAN software (Minitab Inc., State College, PA).

## Results and Discussion

### Stage 1 of the study

#### Nitrogen concentration in agricultural soil

Ten agricultural soils (S1 Table, soils 1-6) were studied for N presence in soil down to 1-meter depth during ordinary agricultural management and the distribution of functional N-related gene. We started by concentrating our work on the analysis of reactive nitrogen content, i.e. ammonium (NH_4_^+^) and nitrate (NO_3_^-^), in soil samples taken every 25 cm in depth, starting from the surface (0 cm) to 1-meter-deep, in the years 2014, 2015 and 2016. Measurements made (*n* = 248) (S5 Table) indicate that despite the soils differing for characteristics and management, with particular reference to N dosed (range of 153-453 kg Ha^-1^ y^-1^) and N-fertilizers used (urea, animal slurries and digestates from animal slurries) (S1 Table), the NO_3_^-^ concentrations (Fig 1) along soil profiles and during agricultural seasons showed similar trends by decreasing dramatically (reduction of 75% ± 14%; *n*=122) proceeding downwards from the soil surface (average of 15.4 ± 11.4 mg N-NO_3_^-^ kg^-1^; *n*=63) to 1 m depth (average of 3.9 ± 4.4 mg N-NO_3_^-^ kg^-1^; *n*=59). Data measured on the surface are similar to those previously reported for agricultural soil [50] but surprisingly, those measured for 75-100 cm depth layer are comparable to the ones reported for natural soil [51]. Ammonium concentration (S2*a* Fig) shows a similar trend, reducing, on average, its content from 4.1 ± 9.6 mg N-NH_4_^+^ kg^-1^ (*n*=63) at the surface (0-25 cm) to 1.6 ± 2.7 mg N-NH_4_^+^ kg^-1^ (*n*=57) at 75-100 cm depth.

**Fig 1.**
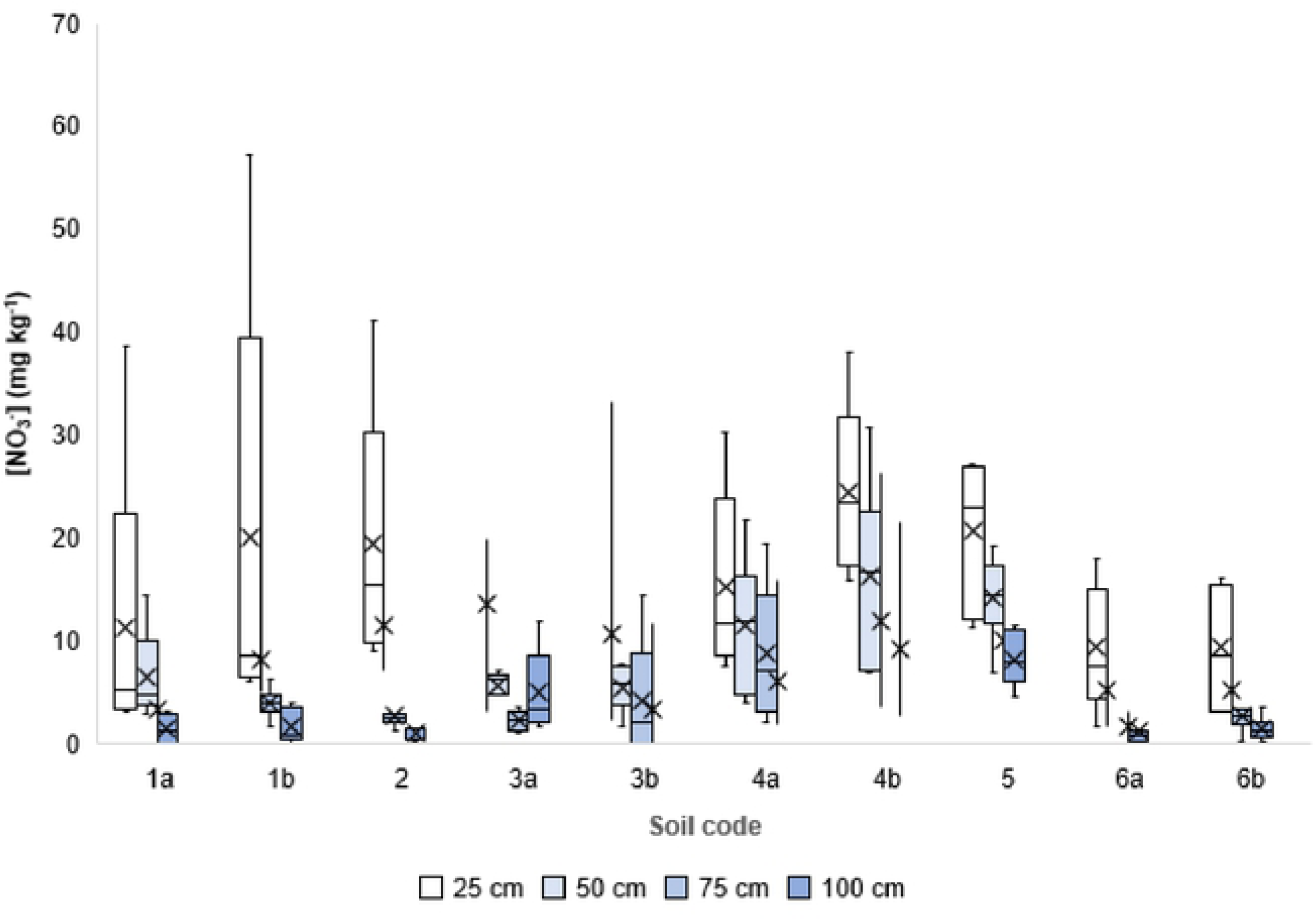
Nitrate concentration in soil. For each experiment, divided for depth classes, the box plot shows minimum and maximum values sampled (bars), the first and the third quartile (boxes), the median (lines inside boxes) and the average (crosses); *n* = 248.

During the agricultural season, NO_3_^-^ content in soils’ surface layers (0-25 cm) varied considerably depending upon: proximity to the fertilization event, total N-dose, presence of crop and sampling date. This trend was not confirmed for the deeper soil layers analysed, that showed much lower variability during the year (Fig 1).

#### The abundance of gene copies related to the N-cycle

Nitrogen transformation in soil is a complex process and depends on many factors, but there is agreement on the fact that that soil microorganisms play a central role [20,22] in regulating the potential for N mineralization, nitrification and denitrification [24]. We measured the DNA gene copies (gene copies g^-1^ soil) coding for enzymes in charge of N nitrification (bacteria genes *amoA*-Eubacteria and *amoA*-Archea), N fixation (bacterial gene *nifH*) and N denitrification (bacterial genes *nirK* and *nosZ*) [52–55], distributed along the profiles of the analysed soils and for all samples taken during the agricultural seasons (*n*=252) (Fig 2 and S6 Table).

**Fig 2.**
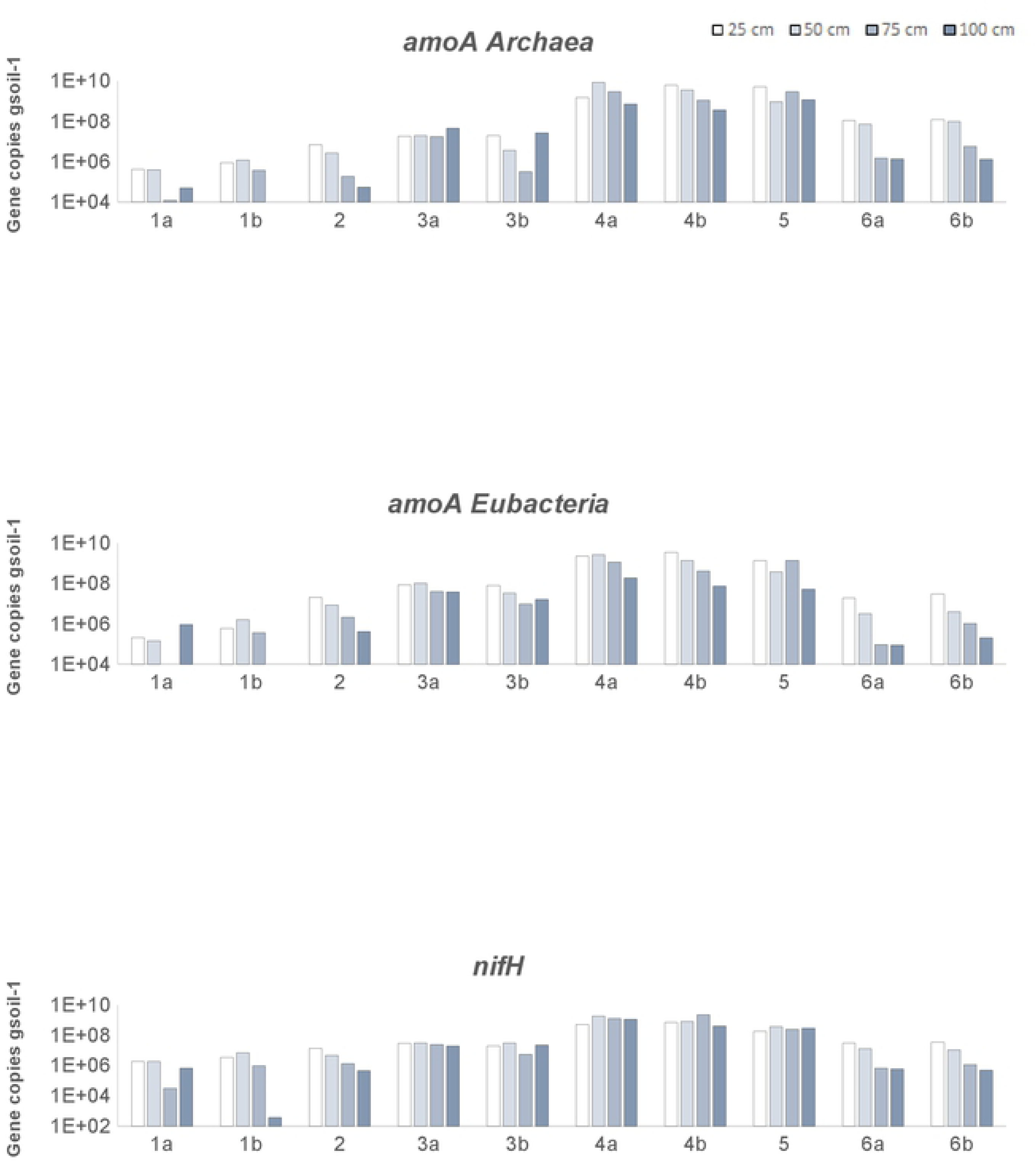
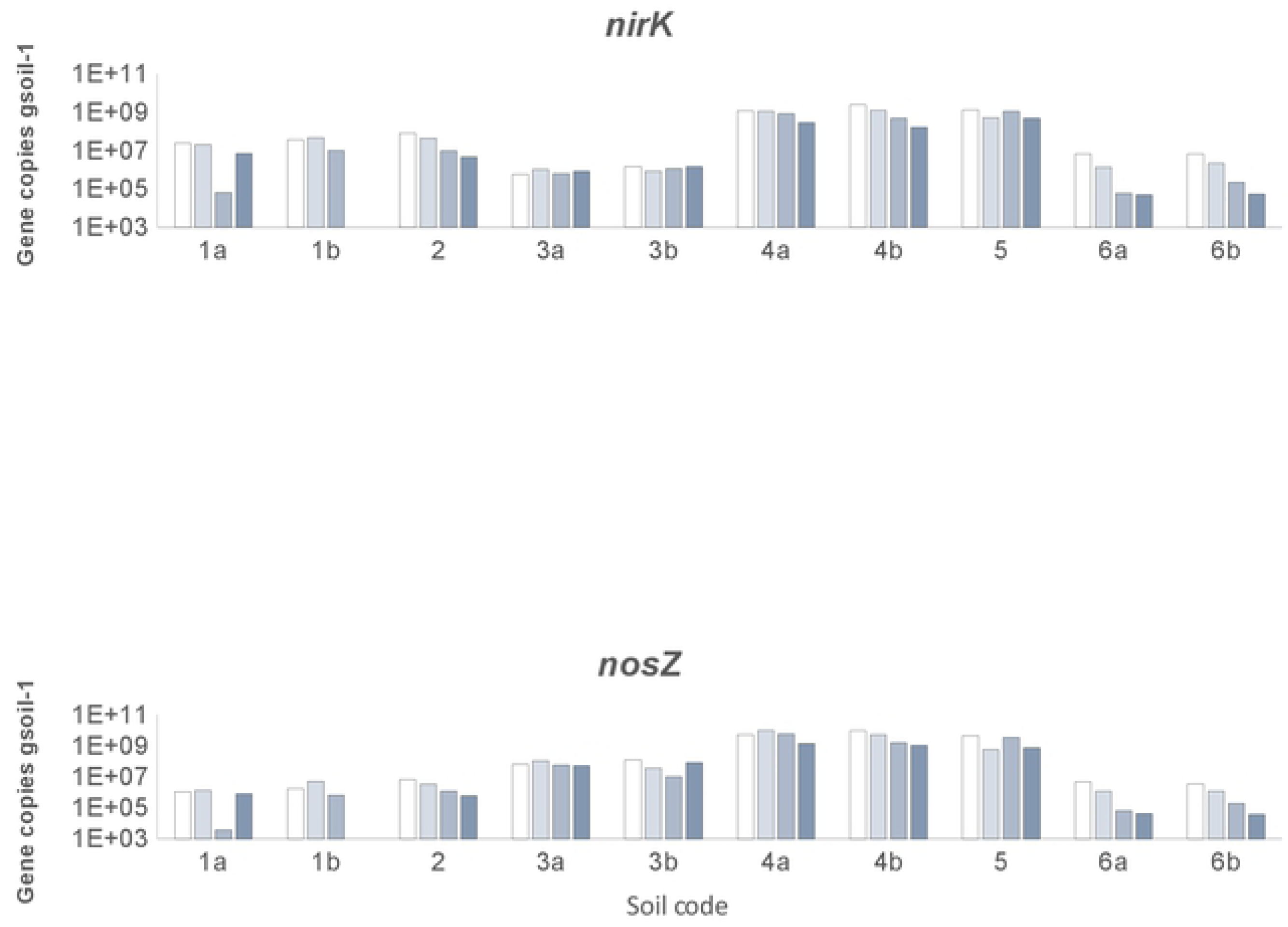
Gene copies concentration in soil. Plots show the number of gene copies (average) for gram of soil for each experiment, grouped by sampling depth. The genes analysed are: amoA from archaea, amoA, nifH, nirK and nosZ from bacteria. Y axis shows log_10_ scale. Error bars show Standard Deviation. n = 252.

Results show a strong spatial coincidence between the number of gene copies detected coding for the different N transformations and mineral nitrogen content (r coefficients > 0.91; *p*<0.05; *n*=252) (S7 Table). These results agree with recent indications highlighting the tendency of soil microorganisms to form complex communities within which nitrogen is metabolized, processed and transformed [22].

Gene copies found per gram of soil decreased with depth for all soils studied (except for the gene *nifH*, related to N fixation) (Fig 3, *p*<0.01; *n*=252).

**Fig 3.**
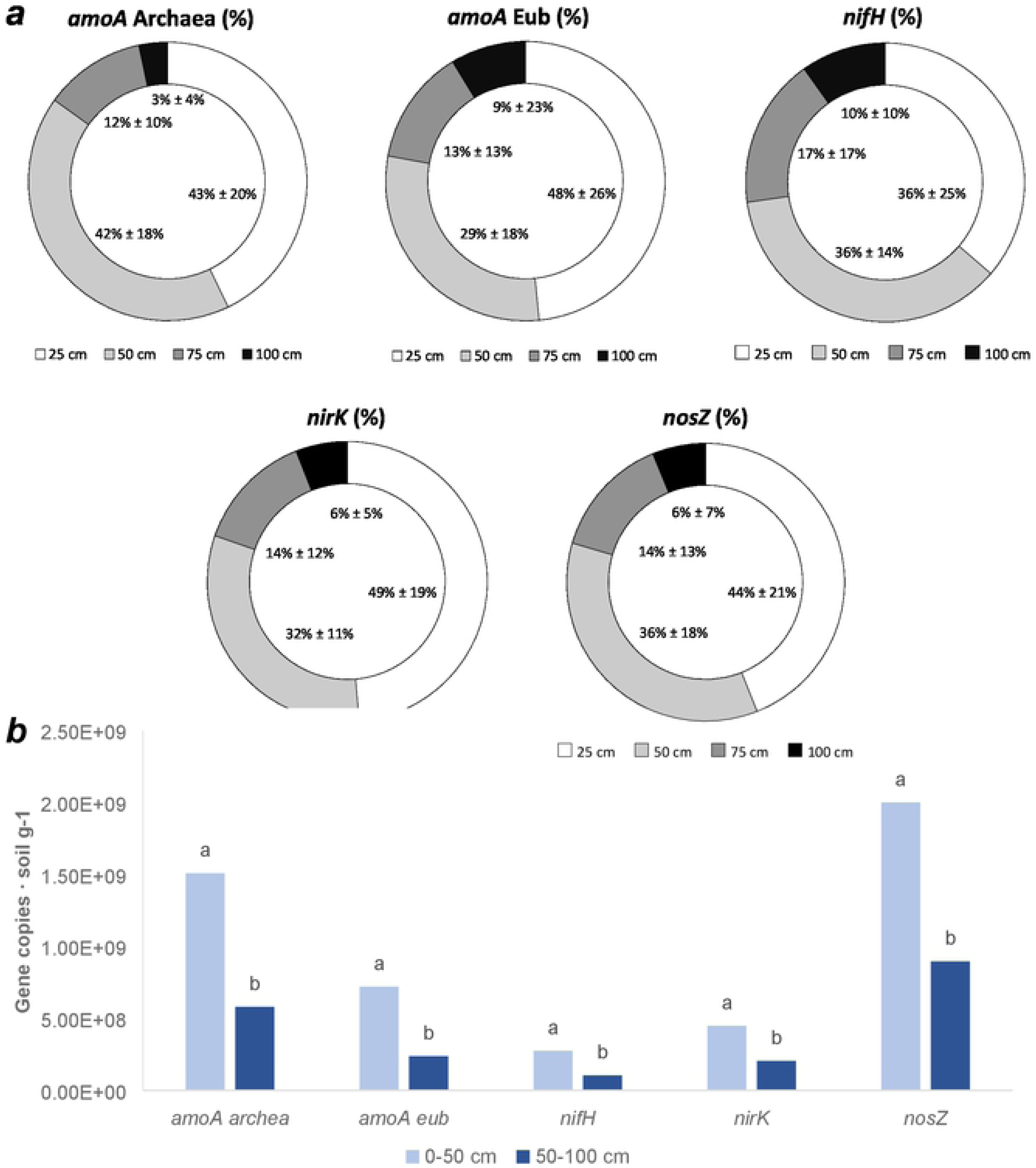
Microorganisms quantification. a: percentages of gene copies in all soils analysed, divided into depth class. Data in the graphic represent the average ± standard deviation of all soils (1a, 1b, 2, 3a, 3b, 4a, 4b, 5, 6a and 6b), (n=252). b: total number of gene copies (average) for gram of soil, divided into two depth classes (0-50 cm; 50-100), (n=252). Letters refer to two-way ANOVA analysis (p<0.01, factor 1: depth (two levels), factor 2: sampling period (in correspondence or not to N fertilization) (two levels) (Gabriel test).

Consequently, we found that the number of genes copies for the N cycle related genes were much higher in the surface layer than in the other layers (Fig 3), in correspondence with the highest concentrations of NO_3_^-^ and NH_4_^+^ (Fig 1). The difference observed was stronger in the case of bacterial amoA genes, i.e. a number of gene copies (gene copies g^-1^ soil) for the layer 75-100 cm, were 16 times lower than those detected in 0-25 cm layer.

### The contribution of the other soil properties to the reduction of nitrate presence at 1-meter depth in soil

Although the abundance of soil microorganisms (i.e. gene copies number) seems to play a fundamental part in determining N speciation in the soils studied, the N dosed, the soil chemical and physical properties, and environmental factors can also play important roles [20,22,56–58]. Therefore, we evaluated the contribution of all these factors to N speciation with particular reference to nitrate by multivariate analysis (principal component analysis, PCA). The two-dimensional PCA graph obtained (Fig 4) shows a clear division of soils into three groups differentiated by NO_3_^-^ concentration at 1-meter depth. In the left quadrants we find soils (group *a*; two-way ANOVA, p=0.0003; F=3.61; DF=18; *n*=59) with very low nitrate concentration at 1-meter depth (1.39 mg N-NO_3_^-^ kg^-1^ ± 1.25; *n*=29) (Fig 4). These soils all showed an alkaline pH (8.45 ± 0.03; *n*= 5) and they are rich in clays and silt (0-50 cm: 28.73% ± 5.07% and 47.14% ± 5.36%, respectively; *n*=5) (S4 Table). These soils showed a strong reduction of the nitrate concentration from the surface to 1 meter of depth (85% ± 19%; *n*=62).

**Fig 4.**
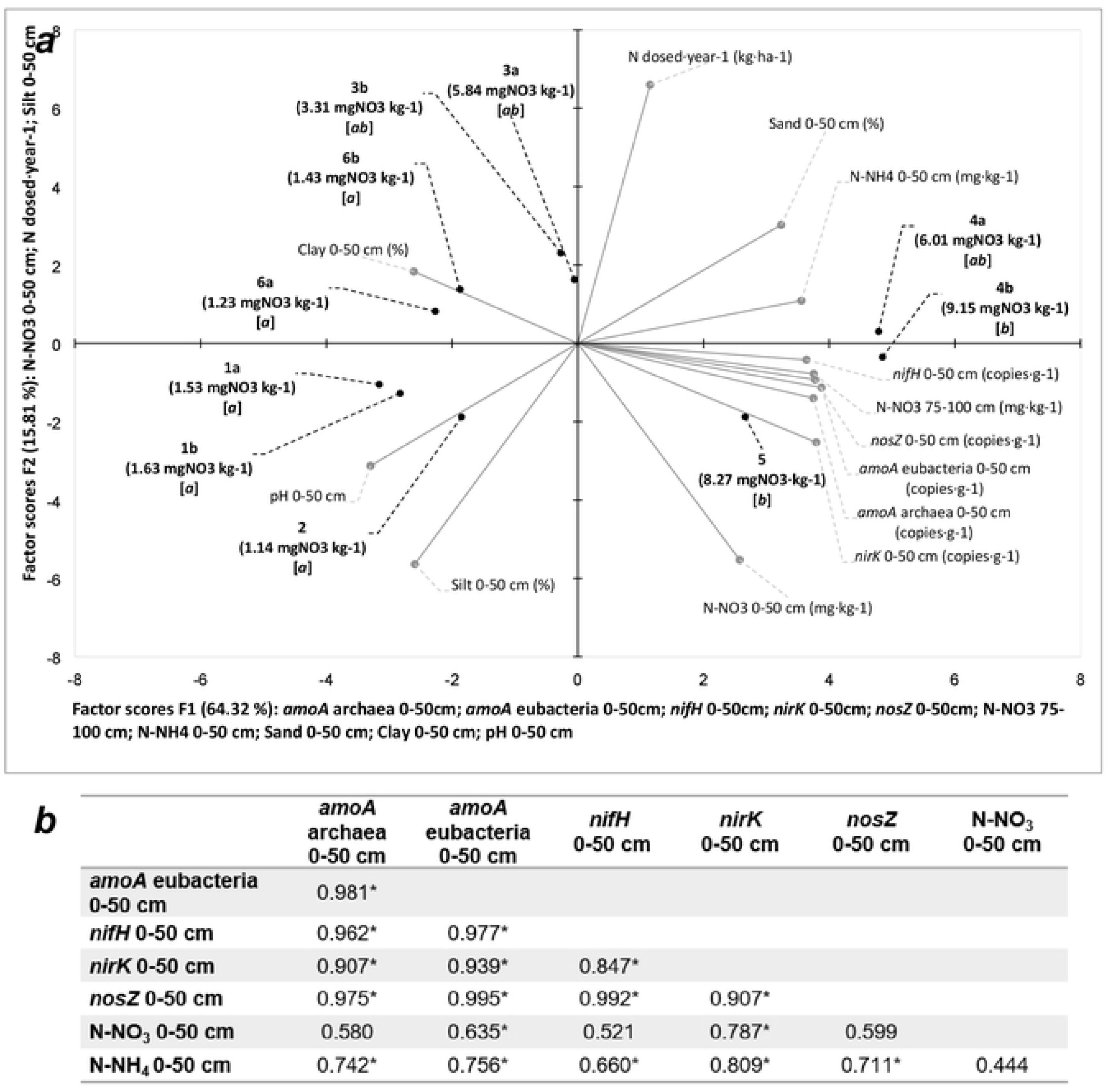
Principal Component Analysis (axes F1 and F2: 80.13 % of total variance). a: PCA output. Grey lines show the projections of the initial variables in the factorial space. Black dots show the position of the experiments in the factorial space. Round brackets show the average concentration of nitrates measured at 1-meter depth. Square brackets show the result of two-way ANOVA analysis on nitrate concentration (p< 0.05; F= 4.1251; Duncan test). Experiments’ labels show the soil code. Axis titles report the variables linked to each axe. **b**: correlation matrix (Pearson) containing: the number of gene copies for gram of soil for amoA archaea, amoA eubacteria, nifH, nirK and nosZ, the N-NO_3_^-^ and N-NH_4_^+^ concentration (mg kg^-1^ of soil, n=189. *= the correlation is significant at level 0.05 (two-tailed test)).

In contrast, the soils which occurred in the right quadrants (group *b*, two-way ANOVA, p=0.0003; F=3.61; DF= 18; *n*=59) (Fig 4) showed a NO_3_^-^ concentration at a 1-meter depth (8.71 mg N-NO_3_^-^ kg^-1^ ± 4.85; *n*=12) significantly greater than that of soils of group a. The pH measured for these soils is neutral (6.93 ± 0.32; *n*=2), and sand is well represented in the first 50 cm (49.95% ± 6.99%; *n*=2) (S4 Table). The nitrate reduction between the surface layer and the 75-100 cm deep layer was less than that measured for the other soil group but still remarkable, i.e. 58% ± 20% (*n*=24). Finally, there is a third group having intermediate values between the other two groups (group *ab*, two-way ANOVA, p=0.0003; F=3.61; DF= 18; *n*=59), i.e. N-NO_3_^-^ of 4.80 mg kg^-1^ ± 4.56; *n*=18 (Fig 4).

Analysis performed (PCA in Fig 4) indicates that, in general, N dose does not affect nitrate presence, unlike the soil texture. In particular clay and sand soil contents seem to affect nitrate concentration along the soil profile (Fig 4), as confirmed by the Pearson correlation analysis for the NO_3_^-^ concentration in the 75-100 cm vs. percentage of sand and clay in the surface layer (0-50 cm), i.e. r=0.718, *p*<0.05, *n*=189 and r=-0.698, *p*<0.05, *n*=189, respectively. pH could also play a role (Fig 4) as it is reported that alkaline pH stimulates biological activities [59], although other authors reported the high adaptability of denitrifying bacteria to different pH [60]. In any case, the reported pH effect contrasts with our results, which indicates that gene copy numbers are higher for those soils characterized by a lower pH (right quadrant of Fig 4) than for soils having alkaline pH (left quadrant of Fig 4). This controversial trend can be explained by considering that gene copies presence is regulated by the amount of reactive N (r > 0.635, *p*<0.05; *n*=189), in agreement with PCA results (Fig 4).

The application of the partial least square analysis (PLS) considering all factors included into PCA analysis gives a regression (R^2^ = 0.96, R^2^cv = 0.95; *p*<0.05; *n*=10; *parameters* = 26) (S8 Table) that confirms all PCA parameters influencing nitrate presence such as before discussed. Indeed, high clay and silt contents reduce nitrate concentration in the 75-100 cm soil layer. On the other hand, soils characterized by light textures (sandy soil) are more exposed to nitrate leaching [34] although in this case an increase in the presence of genes related to nitrifying/denitrifying activities (copies of *amoA*-Eubacteria, *amoA*-Archea, *nirK* and *nosZ* genes) driven by the presence of reactive N in the upper layers (S8 Table), is able to keep nitrate concentration in line with that of natural soils (9.6 mg N-NO_3_^-^ kg^-1^)^35^, emphasizing the primary role of N-cycle related activities in determining the control of nitrate concentration in soils profiles.

Our conclusions in this respect are that nitrate presence at 1-meter soil depth can be explained by the abundance of gene copies for enzymes related to N-cycle, but that soil texture also plays an important role.

### Stage 2 of the study

#### The effect of an excess of N-fertilization on nitrate presence in soil profiles

The results discussed above indicate that soils with different chemical-physical characteristics and treated with different N fertilizers and agronomic N-doses, are able to control nitrate presence in soil profiles, maintaining the NO_3_^-^ concentration detected at one meter depth below that reported for natural (not cultivated) soils at the same depth (10 mg kg^-1^) [51], with NO_3_^-^ (and NH_4_^+^) being cut down, on average, by 75% (and by 71%) passing from the top to the bottom layers.

However, is there a limit beyond which soils’ ability to reduce nitrate presence in 1 m depth soil does end? In order to determine this, we studied nitrate concentration in two soils (Soil 7 and Soil 8) receiving an excess of N fertilization.

Soil 7 showed in autumn (October-November) a high presence of NO_3_^-^ in the surface layer (0-25 cm) (110.3 mg N-NO_3_^-^ kg^-1^ ± 33.6; *n*=6), in correspondence with the high N-fertilization received with pig slurry (S1 Table). Despite this, only a small part of the nitrate reached 75-100 cm depth in soil, the NO_3_^-^ concentration at the same depth in autumn being 6.85 mg N-NO_3_^-^ kg^-1^ ± 1.91 (*n*=6). This value is much lower than that measured in the month of June (15.81 mg N-NO_3_^-^ kg^-1^) after normal N fertilization with urea (138 kg N Ha^-1^) and in the presence of the crop. In this case, differences in nitrate concentration depended on rainfall which was double in June-July in comparison with that for October-November (S9 Table). Soil 8 instead, showed NO_3_^-^ concentrations at the surface which were much higher than those measured for soils fertilized at the normal rate, in particular after high N-fertilization (June and August) (Fig 5c). Nitrate concentration at 1-meter depth in this period was of 32.37 mg N-NO_3_^-^ kg^-1^ ± 23.77 (*n*=3) and in June, the NO_3_^-^ content exceeded 50 mg N-NO_3_^-^ kg^-1^ at 1-meter depth, that is five times higher than values reported (on average) for the soils previously studied, including Soil 7 that was fertilized with an excess of N similarly to Soil 8.

**Fig 5.**
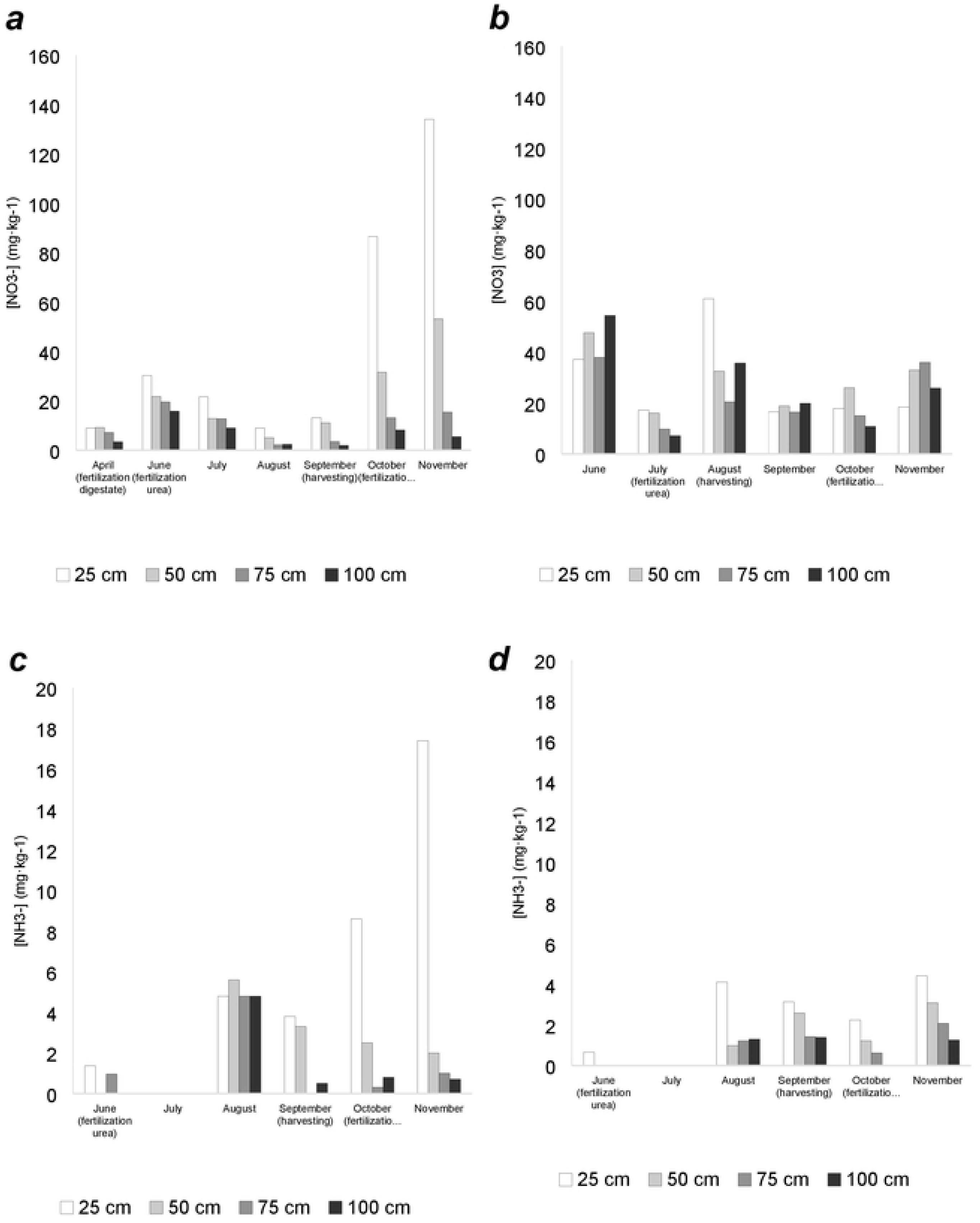
Nitrate (a and b) and ammonium (c and d) concentration in soil of the experiments 7 (a and c) and 8 (b and d) during the year 2016. Experiment 7 was supplied with a total amount of nitrogen of 1,423 kg N Ha^-1^. Experiment 8 was supplied with a total amount of nitrogen of 1,470 kg N Ha^-1^.

It is interesting to compare Soil 7 with Soil 8 in the autumn period. Indeed, in autumn Soil 7 received a large amount of N (860 kg N Ha^-1^) leading to the high NO_3_^-^ presence in the surface layer, which however did not correspond to high nitrate concentration at the 75-100 cm depth soil layer (Fig 5). On the contrary, in the same period Soil 8 received much less N with similar N-fertilizer (580 kg N Ha^-1^) but showed high nitrate presence at 75-100 cm depth, although rainfall registered was much less than that for Soil 7 (205 mm for Soil 7 and 124 mm for Soil 8). This result appears more peculiar if we consider that the number of gene copies g^-1^ related to enzymes implicated in nitrifying-denitrifying activities measured for Soil 8 is of 1 to 2 orders lower than that measured for Soil 7 (Fig 2; S6 Table), despite an alkaline pH (S4 Table) which could stimulate biological activities [59] and an organic carbon content which can support denitrifying activities (S4 Table) [61].

Therefore, both the number of N related gene copies and rainfall events are not consistent with the higher NO_3_^-^ presence at 1-meter depth for Soil 8 and Soil 7, indicating that the reason for the difference that occurs between these two soils should be sought in their own properties [34]. Soil 7 contains more clay, especially in the deeper layers (22 % of clay in the 0-25 cm and 30 % of clay from 25-100 cm) when compared to Soil 8 (from 17.4 % in the upper layer to 10.4 % in the deeper layer) and we regard this difference as being able to explain the different results on nitrate presence [20]. We can therefore summarize results as follows: Soil 7 received a large amount of ammonia in autumn that was transformed into nitrate by ammonia oxidation microorganisms, then nitrate was concentrated above all in the surface layer, because it was not rapidly leached by rainwater. In this way, the high residential time of ammonia and nitrate due to the abundant clay presence in the soil allowed its denitrification, explaining the low NO_3_^-^ content found at 1-meter depth [62]. This fact was confirmed by both the higher number of gene copies related to nitrifying/denitrifying activities registered for Soil 7 with respect to Soil 8 (S6 Table), and by the very high positive correlation found for genes coding for nitrification with those for denitrification (*amoA* Archaea vs *nirK*: r = 0.814, *p*<0.05; *amoA* EUB vs *nirK*: r = 0.991, *p*<0.01; *n*=8; layer 0-25 cm). The fact that the number of gene copies did not decrease along the soil profile seems to indicate that nitrifying-denitrifying processes continue throughout all soil depths (S6 Table). Contrarily, Soil 8, due to the high presence of sand, led to the rapid leaching of nitrate, limiting the proliferation of microorganisms (*i.e.* gene copies number) related to N denitrification, that are, in effect, much lower than those measured for Soil 7 (S6 Table).

## Conclusions

Results of this work suggest that with a normal N fertilization (up to 450 kg N Ha^-1^) the microbial populations of the soil involved in the N cycle were able to completely metabolize the nitrogen supplied with fertilization, despite soil characteristics, ensuring low nitrate content at one-meter depth. However, for higher N fertilization rates (1,243 kg N Ha^-1^ and 1,470 kg N Ha^-1^), the activity of soil microorganisms was not able to metabolize all the nitrogen. In this case the characteristics of the soil, i.e. texture, and rain precipitation, also regulated the presence of nitrate in soil profiles.

## Author Contributions

**Conceptualization:** Fabrizio Adani

**Data curation**: Fabrizio Adani, Massimo Zilio

**Formal analysis:** Barbara Scaglia, Massimo Zilio, Fabrizio Adani

**Funding acquisition:** Gabriele Boccasile, Fabrizio Adani

**Investigation:** Fabrizio Adani, Silvia Motta, Fulvia Tambone, Andrea Squartini

**Methodology:** Fabrizio Adani, Massimo Zilio, Andrea Squartini

**Project Administration:** Fabrizio Adani

**Resources:** Fabrizio Adani, Silvia Motta, Fulvia Tambone, Andrea Squartini

**Supervision**: Fabrizio Adani

**Validation:** Massimo Zilio, Fabrizio Adani

**Visualization**: Massimo Zilio

**Writing – Original Draft Preparation:** Massimo Zilio, Fabrizio Adani

**Writing – Review & Editing:** Massimo Zilio, Fabrizio Adani

## Acknowledgment

CONVENZIONE QUADRO Regione Lombardia – ERSAF, Italy: Supporto tecnico per l’applicazione e il monitoraggio della direttiva nitrati (ARMOSA) + rapporto ambientale per VAS, piano triennale 2014 – 2016, Agreement Collaborazione tecnico-scientifica per azioni finalizzate a valutare la sostenibilità complessiva della gestione”. Project N. 15-3-3014000-218 and 15-3-3014000-219. Gruppo Ricicla lab. Università degli Studi di Milano, Italy, Project N. 1705, 2011: RV_ATT_COM16FADAN_M.

## Supporting information

**S1 Fig. Geographical location of the soils exploited in this work.** The reported cartographic table, excluding the points of interest, was obtained from OpenStreetMap.org. The data is available under the Open Database License, and the cartographic table is published with a CC BY-SA license.

**S2 Fig. Ammonium (*a*) and phosphate (*b*) concentration in soil.** For each experiment, divided for depth classes, the box plot shows minimum and maximum values (bars), the first and the third quartile (boxes), the median (lines inside boxes) and the average (crosses); *n*=248.

**S1 Table. Soil studied and soil agronomic management.** In the Table are reported the agronomic data of all the experiments made.

**S2 *Table*. Sampling periods.** The months of the year in which at least one sampling was carried out for the corresponding Soil are marked in grey.

**S3 Table. Primers used in the quantification of nitrogen related microorganisms in soil.** For each primer are shown name, nucleotide sequence, target gene and reference. N = undefined nucleotide.

**S4 Table. Soil characteristics.** Texture (sand tot. silt tot and clay) and pH (H_2_O) for the four soil depths studied. USDA soil taxonomy, percentage of organic carbon (OC%) and cation exchange capacity (CEC) for the 0-25 cm layer.

**S5 Table. Nitrate, ammonium.** Their concentration (average ± standard deviation) in the analysed soils.

**S6 Table. Gene copies for every microorganism class in the soils analysed**. Average ± standard deviation

**S7 Table. Pearson correlation matrix.** The test was performed using the data from all the regularly fertilized farms (1-6). Numbers in table indicate the r correlation coefficients.

**S8 Table. PLS regression.** Regression for nitrate content in soils (75-100 cm) (*n* = 10) vs. agronomical, chemical, physical, biological and meteorological data (parameters = 26). In the table are reported variables included into regression and their importance.

**S9 Table. Rainfall**. Total mm of water from rainfall during the year for every soil (2014, 2015 or 2016), divided for months.

